# A mathematical model of calcium dynamics: Obesity and mitochondria-associated ER membranes

**DOI:** 10.1101/477968

**Authors:** Jung Min Han, Vipul Periwal

## Abstract

Multiple cellular organelles tightly orchestrate intracellular calcium (Ca^2+^) dynamics to regulate cellular activities and maintain homeostasis. The interplay between the endoplasmic reticulum (ER), a major store of intracellular Ca^2+^, and mitochondria, an important source of adenosine triphosphate (ATP), has been the subject of much research, as their dysfunctionality has been linked with metabolic diseases. Interestingly, through out the cell’s cytosolic domain, these two organelles share common microdomains called mitochondria-associated ER membranes (MAMs), where their membranes are in close apposition. The role of MAMs is critical for intracellular Ca^2+^ dynamics as they provide hubs for direct Ca^2+^ exchange between the organelles. A recent experimental study reported correlation between obesity and MAM formation in mouse liver cells, and obesity-related cellular changes that are closely associated with the regulation of Ca^2+^ dynamics. We constructed a mathematical model to study the effects of MAM Ca^2+^ dynamics on global Ca^2+^ activities. Through a series of model simulations, we investigated cellular mechanisms underlying the altered Ca^2+^ dynamics in the cells under obesity. We found that the formation of MAMs is negatively correlated with the amplitude of cytosolic Ca^2+^ activities, but positively correlated with that of mitochondrial Ca^2+^ dynamics and the overall frequency of Ca^2+^ oscillations. We predict that, as the dosage of stimulus gradually increases, liver cells from obese mice will reach the state of saturated cytosolic Ca^2+^ concentration at a lower stimulus concentration, compared to cells from healthy mice.

**Author summary:** It is well known that intracellular Ca^2+^ oscillations carry encoded signals in their amplitude and frequency to regulate various cellular processes, and accumulating evidence supports the importance of the interplay between the ER and mitochondria in cellular Ca^2+^ homeostasis. Miscommunications between the organelles may be involved in the development of metabolic diseases. Based on a recent experimental study that spotlighted a correlation between obesity and physical interactions of the ER and mitochondria in mouse hepatic cells, we constructed a mathematical model as a probing tool that can be used to computationally investigate the effects of the cellular changes linked with obesity on global cellular Ca^2+^ dynamics. Our model successfully reproduced the experimental study that observed a positive correlation between the ER-mitochondrial junctions and the magnitude of mitochondrial Ca^2+^ responses. We postulate that hepatic cells from lean animals exhibit Ca^2+^ oscillations that are more robust under higher concentrations of stimulus, compared to cells from obese animals.

## Introduction

In most multicellular organisms, calcium (Ca^2+^) is a ubiquitous second messenger that controls a vast array of cellular activities spanning from cell birth to apoptosis [1]. The endoplasmic/sarcoplasmic reticulum (ER/SR) and mitochondria have been the center of attention in the study of intracellular Ca^2+^ dynamics, due to their role as internal Ca^2+^ stores. The SR is mostly found in muscle cells, which are not the subject of this paper, so we only refer to the ER. It has been suggested that dysfunction of Ca^2+^ regulation in the ER and/or mitochondria leads to disrupted cellular homeostasis, and is associated with pathological processes, including metabolic diseases and neurodegenerative diseases [2–7].

Upon agonist stimulation, almost all types of cells exhibit fluctuations in cytosolic Ca^2+^ concentration, phenomena often referred to as Ca^2+^ oscillations, with signals encoded in oscillation frequencies and amplitudes. Among many cellular compartments, the ER, whose internal Ca^2+^ concentration is three to four orders of magnitude larger than that of the cytosol in resting condition, is considered as the main contributor to the generation of Ca^2+^ oscillations. The ER has several types of Ca^2+^ channels on the membrane that release Ca^2+^ once activated. The most well-studied Ca^2+^ release channels are inositol trisphosphate receptors (IPRs) and ryanodine receptors (RyRs). As a high cytosolic Ca^2+^ concentration is toxic and often leads to cell death, released Ca^2+^ is quickly pumped back into the ER lumen through sarco/endoplasmic reticulum Ca^2+^ ATPase (SERCA) pumps, which consume energy to sequester Ca^2+^ against its concentration gradient. Some Ca^2+^ released from the ER can be taken up by mitochondria through the mitochondrial Ca^2+^ uniporter (MCU), and then released back to the cytosol via the sodium/calcium exchanger (NCX). Thus, it is generally accepted that mitochondria have the ability to modulate oscillation frequencies and amplitudes, and consequently, affect the progression of cellular activities [4].

Having a spatially extended membrane network, the ER is often positioned in close proximity with other cellular organelles and forms membrane contact sites. Such sites between the ER and mitochondria are called mitochondria-associated ER membranes (MAMs), and it has been suggested that they play a critical role in Ca^2+^ exchange between the organelles [4, 5]. Since mitochondrial Ca^2+^ regulation is closely linked with adenosine triphosphate (ATP) synthesis and reactive oxygen species (ROS) production [8], understanding the mechanisms underlying the ER-mitochondrial Ca^2+^ crosstalk is of great scientific and physiological interest. A major advantage of MAM formation is that due to its minuscule size, even a small Ca^2+^ flux into the domain would be amplified, which is convenient for the MCUs which have a low Ca^2+^ affinity, i.e., they require a high concentration of Ca^2+^ in order to activate.

Arruda et al. [9] reported a positive correlation between obesity and the degree of MAM formation. They also found different expression levels of Ca^2+^ channels between liver cells of lean and obese mice. These findings indicate the possibility of obesity-induced changes in Ca^2+^ dynamics in MAMs, and consequently, in the ER as well as mitochondria. Indeed, liver cells from obese animals showed higher baselines of cytosolic Ca^2+^ concentration and mitochondrial Ca^2+^ concentration, compared to cells from lean mice. Furthermore, Ca^2+^ transients generated from ATP stimulation led to higher concentration peaks in obese mouse mitochondria. Interestingly, this observation was not accompanied by higher peaks in cytosolic Ca^2+^ concentration, i.e., cells from obese and lean mice exhibited similar ATP-induced rises in cytosolic Ca^2+^ concentration.

Computational models of experimental data have been a valuable tool for understanding the dynamics of intracellular Ca^2+^. Most models have focused on either the ER Ca^2+^ handling [10–13] or mitochondrial Ca^2+^ dynamics [14–17] and only a handful of them have integrated the dynamics from both organelles [18, 19]. Recently, there have been an increasing number of studies that combined both experimental and theoretical approaches to probe the cellular mechanisms underlying Ca^2+^ crosstalk between the ER and mitochondria. The model proposed by Szopa et al. [20] assumes that due to the minuscule volume of MAMs, the MCUs in MAMs sense Ca^2+^ concentration in the ER. Thus, the MCU Ca^2+^ flux in their model is essentially direct Ca^2+^ flow from the ER. Using numerical methods, they investigated the effects of this flow on the shape (bursting) and period of Ca^2+^ oscillations, and observed that mitochondrial Ca^2+^ concentrations tend to a high level in some regions of parameter space. Another recent model by Qi et al. [21] considers a range of possible distances between the IPRs and MCUs in MAMs, and expresses Ca^2+^ concentration in MAMs as a solution to a linearized reaction-diffusion equation. In this model, the concentration of Ca^2+^ that is sensed by the MCUs in MAMs depends on the distance of the MCUs from the point of source (a cluster of IPRs) and how fast Ca^2+^ diffuses in MAMs. The authors showed that Ca^2+^ signals can be significantly modulated by the distance, and determined an optimal distance between the IPRs and MCUs for effective Ca^2+^ exchange for the generation of Ca^2+^ oscillations. On the other hand, Wacquier et al. [22] published a model that associates Ca^2+^ oscillations with mitochondrial metabolism, and investigated the role of mitochondrial Ca^2+^ fluxes on the oscillation frequency. Their model modified one of the parameters that describes the Ca^2+^ concentration for the activation of the MCUs to a lower concentration than the one originally suggested by Magnus and Keizer [16]. By doing so, they implicitly included MAMs, with the following assumption: MCUs are activated at the average concentration of Ca^2+^ in the whole cytosol (including MAMs). They found that mitochondrial Ca^2+^ fluxes can modulate the frequency of Ca^2+^ oscillations.

Here, we construct a mathematical model to investigate the cellular mechanisms underlying the altered mitochondrial Ca^2+^ dynamics observed in obese mice. The model extends the model of Wacquier et al. [22], and explicitly includes Ca^2+^ dynamics in MAMs. We incorporated the model structure proposed by Penny et al. [23], wherein the cytosol is compartmentalized to two separate domains: the bulk cytosol and membrane contact sites between organelles. Rather than expressing the Ca^2+^ concentration in MAMs as an algebraic function of that of the cytosol, we model it as a dynamic variable that is determined by influxes and effluxes of the domain. We investigated how Ca^2+^ signals are affected by the obesity related changes in Ca^2+^ channel expression levels.

## Materials and Models

A better way of understanding the model structure is to consider the model as two sub-models that are coupled by common factors, Ca^2+^ and ATP. One of the sub-models describes intracellular Ca^2+^ dynamics, while the other models mitochondrial metabolic pathways and voltage.

### Ca^2+^ dynamics

Jones et al. [24] observed vasopressin-induced Ca^2+^ oscillations in hepatocytes in Ca^2+^-free environment. This experiment suggested that hepatocytes do not require Ca^2+^ influx from the extracellular space to trigger Ca^2+^ oscillations, and that their oscillations are primarily caused by the periodic release and re-uptake of Ca^2+^ by intracellular stores. During the experiment, the oscillations were sustained for about 15 minutes after the stimulation, with decreasing frequencies, most likely due to the continuous loss of Ca^2+^ across the plasma membrane as, eventually, cells became completely depleted of Ca^2+^. When modeling Ca^2+^ signals with these behaviors, it is useful to consider a closed-cell model, where it is assumed that the total intracellular Ca^2+^ is fixed (i.e., a cell does not lose or gain Ca^2+^ through the plasma membrane). Since the main focus of this study is on Ca^2+^ dynamics in liver cells, we approached the problem with a closed-cell model. We first compartmentalized the cellular domain into four separate regions: the ER, the bulk cytosol, a mitochondrion, and MAM, and assumed that Ca^2+^ concentration within each region, denoted by *C_ER_*, *C_cyt_*, *C_mito_*, and *C_MAM_*, respectively, is determined by Ca^2+^ influxes and effluxes going in and out of that region. Fig. 1 shows a schematic diagram of the compartments and Ca^2+^ fluxes in the model. Since the model does not include Ca^2+^ fluxes across the plasma membrane, the total intracellular Ca^2+^ concentration, *C*_t_, is written as:

**Fig 1.**
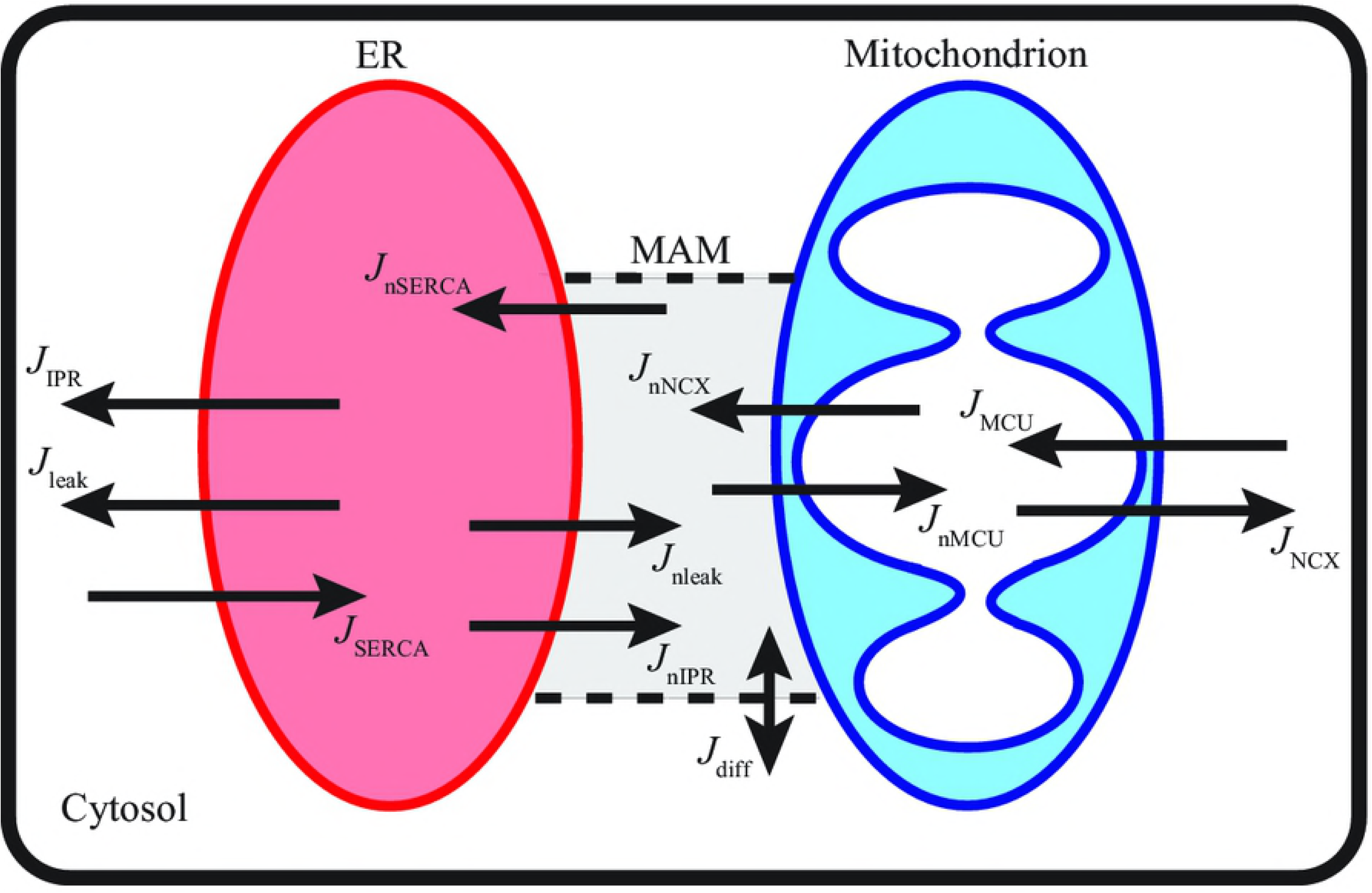
Schematic diagram of Ca^2+^ dynamics described by the model. The ER releases Ca^2+^ to the cytosol and MAM; the fluxes are denoted as *J*_IPR_ and *J*_nIPR_, respectively. The cytosolic Ca^2+^ is pumped back into the ER lumen via SERCA (*J*_SERCA_), and the Ca^2+^ in MAM are also pumped back into the ER (*J*_nSERCA_). Mitochondria uptake Ca^2+^ from the cytosol (*J*_MCU_) and MAM (*J*_nMCU_), and mitochondrial Ca^2+^ is exchanged with Na^+^ in the cytosol (*J*_NCX_) and MAM (*J*_nNCX_). Ca^2+^ can freely diffuse between the cytosol and MAM (*J*_diff_).

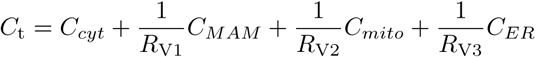

The *R*_V_’s account for compartment volume differences and are defined as:

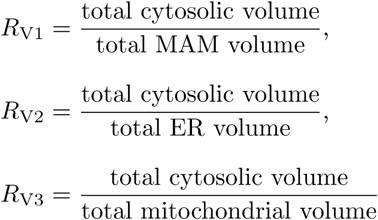

Given that the mitochondrial outer membrane is freely permeable to small molecules, such as Ca^2+^, through the voltage dependent anion channel (VDAC), we assume that Ca^2+^ concentration in the inter-membrane space is equivalent to that of the cytosol. Thus, there is one effective layer of impermeable boundary, the mitochondrial inner membrane, that separates cytosolic Ca^2+^ from mitochondrial Ca^2+^.

The Ca^2+^ dynamics part of the model consists of the following ordinary differential equations (ODEs):

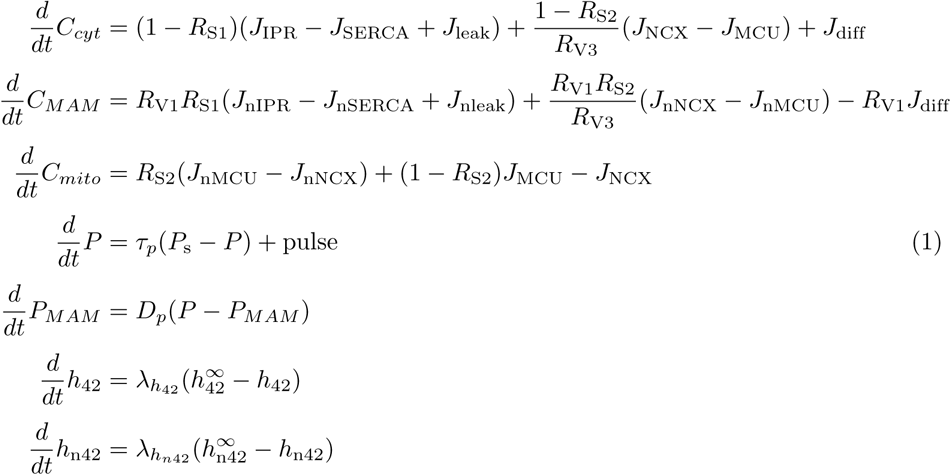

*P* and *P*_*MAM*_ represent the concentrations of IP_3_ in the bulk cytosol and the MAM, respectively. *h*_42_ and *h*_n42_ denote the activation variables of the IPRs in the bulk cytosol and the MAM, respectively. The *R*_S_’s are surface ratios, defined as

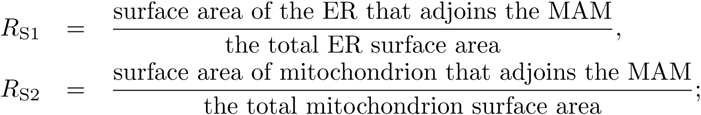

see Table for their values. Short descriptions for the *J_*_* Ca^2+^ fluxes are given below.

- *J*_IPR_ and *J*_nIPR_: IPR Ca^2+^ flux into the bulk cytosol and MAMs from the ER
- *J*_SERCA_ and *J*_nSERCA_: SERCA pump Ca^2+^ flux into the ER from the bulk cytosol and MAMs
- *J*_leak_ and *J*_nleak_: a small Ca^2+^ leak from the ER into the bulk cytosol and MAMs
- *J*_NCX_ and *J*_nNCX_: NCX Ca^2+^ flux from mitochondria into the bulk cytosol and MAMs
- *J*_MCU_ and *J*_nMCU_: MCU Ca^2+^ flux into mitochondria from the bulk cytosol and MAMs
- *J*_diff_ : Ca^2+^ diffusion between the bulk cytosol and MAMs

## IPR model

We incorporated the IPR model proposed in Cao et al. [25], which assumes that the receptors are either in drive mode when they are mostly open, or in park mode when they are mostly closed. The drive mode has one open state (*O*_6_) and one closed state (*C*_2_), while there is one closed state (*C*_4_) in the park mode. The transition rates between the modes are denoted by *q*_24_ and *q*_42_, and the rates between the states within the drive mode are *q*_26_ and *q*_62_. The open probability of the drive mode is *q*_26_*/*(*q*_26_ + *q*_62_) (*≈* 70%). *q*_24_ and *q*_42_ are given by

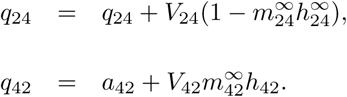

The *m*’s and *h*’s are gating variables that govern the opening and closing kinetics of the receptors, with the following quasi-equilibria:

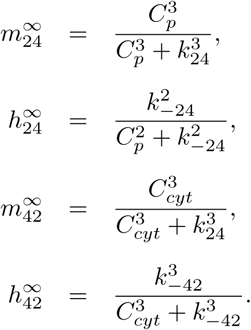

While *m*_24_, *m*_42_, and *h*_24_ are assumed to have reached their quasi-equilibria instantaneously, there is the rate at which *h*_42_ approaches its equilibrium, 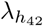,

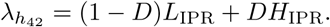

*C*_*p*_ = *C*_*p*0_(*C*_*ER*_/680) denotes the concentration of Ca^2+^ at the pore of a receptor. The expressions for the *V*’s, *a*’s, and *k*’s are

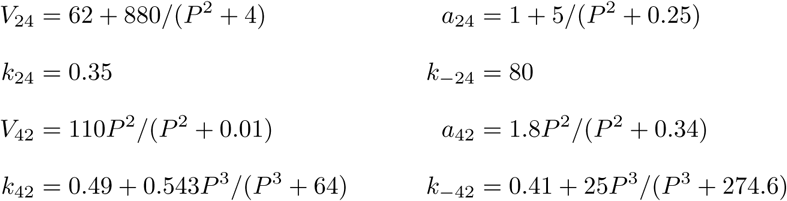

The open probability of the IPRs in the cytosol, *O*_IPR_ is defined as

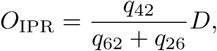

where *D*,

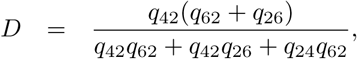

represents the proportion of the IPRs that are in the drive mode. Then the IPR fluxes are modeled as:

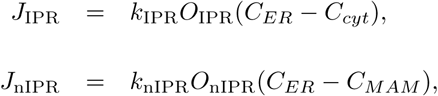

where *O*_nIPR_ is in the same functional form as *O*_IPR_, except that the composing functions are now expressed with *C_MAM_* and *P*_*MAM*_, instead of *C_cyt_* and *P*.

The diffusion flux between the MAM and the cytosol is a linear function of the concentration difference,

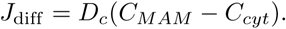

Small leak fluxes across the ER membrane are given by

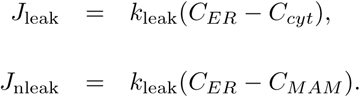

We follow Wacquier et al. [22] in modeling the SERCA, the MCU, and the NCX fluxes,

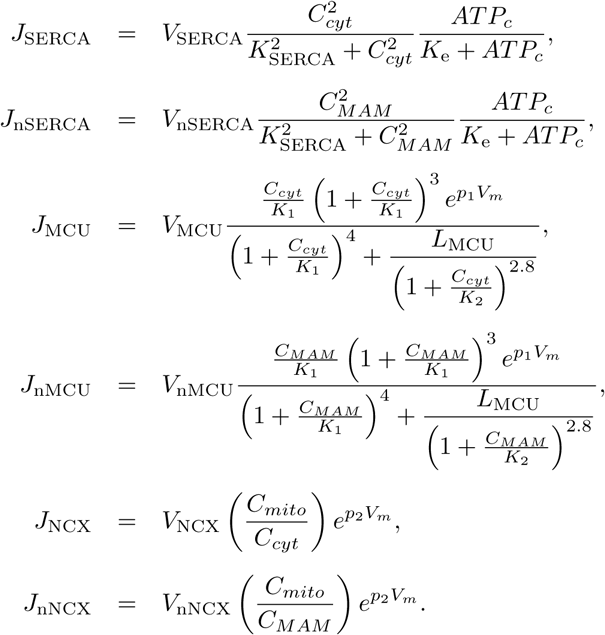

They set the parameter *K*_1_ to 6 *µ*M, but this represents the average level of Ca^2+^ in the whole cytosol, including MAMs, when the concentration reaches a physiologically reasonable level in MAMs. Since we consider MAMs explicitly in the model, we set *K*_1_ to 19 *µ*M, as originally proposed by Magnus et al. [16].

### IP_3_ metabolism

A number of studies suggest Ca^2+^ oscillations in hepatocytes can occur at a constant level of IP_3_. In particular, an experimental study reported oscillating Ca^2+^ concentrations in the ER lumen in permeabilized hepatocytes, while the concentration of IP_3_ was clamped at a submaximal concentration [26]. Thus, we assumed that Ca^2+^ oscillations are primarily generated by Ca^2+^ feedback on the opening and closing kinetics of the IPR. For simplicity, we did not consider cellular formation or breakdown of IP_3_ that involves Ca^2+^ or protein kinase C (PKC). Instead, we model IP_3_ dynamics as a gradual increase from 0 *µ*M to its steady-state concentration, *P*_s_, at a rate of *τ_p_*. However, if hepatocytes were to exhibit IP_3_ oscillations, it is possible to include positive and/or negative Ca^2+^ feedback on IP_3_ metabolism in the model to introduce IP_3_ oscillations as passive reflections of Ca^2+^ oscillations.

Apart from continuous stimulation, the model can be perturbed with a single stimulation in a pulsatile manner. In such a case, cells would be exposed to a certain amount of agonist for a short period of time. While continuous stimulation saturates the concentration of IP_3_ to its steady state, a pulse stimulation produces a sudden increase in IP_3_ concentration, followed by a natural decay at a rate of degradation. In the model, a pulse of stimulation is described by

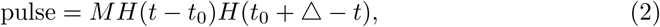

where *H* is the Heaviside function

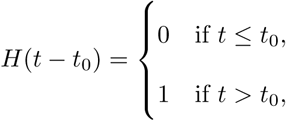

*M* is the pulse magnitude, *t*_0_ is the time at which the pulse is given, and *Δ* is the pulse duration.

### Mitochondrial metabolic pathway model

The main function of mitochondria is to create ATP by oxidative phosphorylation. Due to this particular role, mitochondria are the powerhouse of the cell. The mitochondrial metabolic pathway is initiated by the uptake of pyruvate, which is the end product of cytosolic glycolysis. Pyruvate in the mitochondrial matrix then enters the tricarboxylic acid (TCA) cycle, also known as the citric acid cycle or the Krebs cycle, to generate the reducing agent NADH that has electrons with a high transfer potential. The concentration of mitochondrial NADH can also be increased by the activity of the malate-aspartate shuttle (MAS). NADH then goes through an electron transport chain (ETC), where the electrons are separated and used to drive protons (H^+^) across the inner membrane and generate a proton gradient between the intermembrane space and mitochondrial matrix. As protons accumulate in the intermembrane space, the gradient across the inner membrane is used by the F1FO-ATPase to convert mitochondrial adenosine diphosphate (ADP) to ATP via phosphorylation. The produced ATP is then transported to the cytosol by adenine nucleotide translocases (ANT), which carry out the exchange of cytosolic ADP and mitochondrial ATP across the inner mitochondrial membrane.

Ca^2+^ is an important component in mitochondrial metabolism, as it promotes the production of NADH. An increase in mitochondrial Ca^2+^ concentration upregulates the TCA cycle, and an increase in cytosolic Ca^2+^ concentration stimulates the aspartate-glutamate carrier (AGC), a protein involved in the MAS. We incorporated a part of the model in Wacquier et al. [22] to describe Ca^2+^-stimulated mitochondrial metabolism. We combined the calcium model with a model of mitochondrial metabolic pathways proposed by Wacquier et al. [22]:

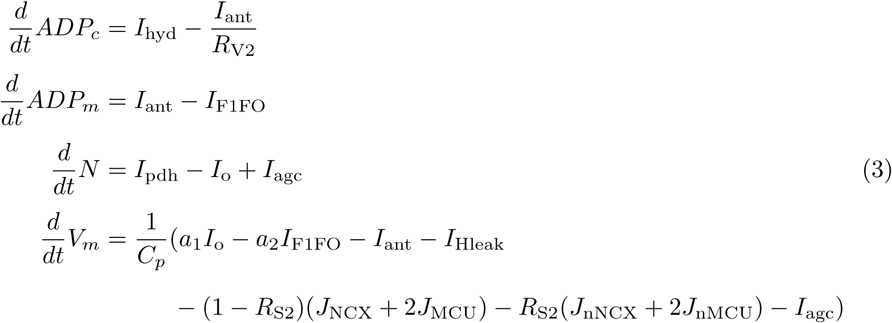

The variables *ADP*_*c*_ and *ADP*_*m*_ measure ADP concentrations in the cytosol and mitochondrion, while *N* is the concentration of mitochondrial NADH. *V*_*m*_ models the voltage difference across the inner mitochondrial membrane. The *I*_*\**_ rates are:

- *I*_hyd_: rate of ATP hydrolysis
- *I*_ant_: rate of the ADP/ATP translocator
- *I*_F1FO_: rate of ADP phosphorylation
- *I*_pdh_: the production rate of NADH by the pyruvate dehydrogenase
- *I*_o_: rate of NADH oxidation
- *I*_agc_: the production rate of NADH from the MAS
- *I*_Hleak_: the ohmic mitochondrial proton leak

Along with the conservation of the total intracellular Ca^2+^ concentration, *C*_t_, the model suggests the conservation of the following ion concentrations: total NADH (oxidized and reduced), mitochondrial di- and trisphosphorylated adenine nucleotides, and cytosolic di- and trisphosphorylated adenine nucleotides. Mathematically speaking,

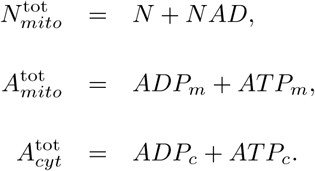

Other functions of the mitochondrial model are reproduced below for convenience. The model parameters are in Table. For modeling purposes, some of the parameters are modified from their original values as in Wacquier et al. [22]. We find these modifications justifiable, as the original values were chosen by the authors to reproduce their experimental data, and hence were not based on any physiological evidence.

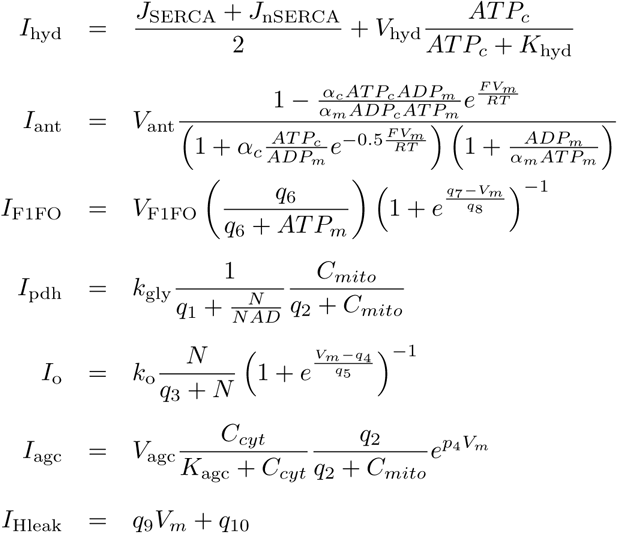

### Numerical simulations

All the numerical simulations presented in this paper were computed with XPPAUT [27].

## Results

### Up-regulation of MAMs and mitochondrial Ca^2+^ activities

Due to the morphology of MAMs, where the ER and mitochondria are in close contact, it is generally accepted that there is direct Ca^2+^ exchange between the organelles at such sites. Based on this notion, it is reasonable to expect that an increase in the degree of MAM formation up-regulates mitochondrial Ca^2+^ intake via MAMs, and consequently induces larger amplitudes of mitochondrial Ca^2+^ signals. In fact, Arruda et al. [9] used synthetic linkers to mechanically induced more MAMs in hepatocyte-derived mouse Hepa1-6 cells, and observed higher ATP-stimulated mitochondrial Ca^2+^ peaks, compared to the cells without the linkers. This experiment revealed a correlation between mitochondrial Ca^2+^ activity and ER–mitochondrial interactions. Through the following model simulations, we confirmed that the model behaves in this manner.

To determine whether the model can reproduce the experimental observations of Arruda et al. [9], we performed model simulations using two different sets of values for the *R*_S_ parameters, the portions of the ER and mitochondrial membranes that face each other. The model with (*R*_S1_*, R*_S2_) = (0.2, 0.1) was stimulated with a pulse of IP_3_, generated from Eq. 2 with *M* = 0.5, *t*_0_ = 4, and Δ = 0.5. Then the same stimulation was applied to the model with (*R*_S1_*, R*_S2_) = (0.3, 0.15), which mimics the up-regulation of MAMs induced by the synthetic linkers. The cytosolic Ca^2+^ transient was not affected by the parameter increases, however, the mitochondrial Ca^2+^ transient exhibited a larger amplitude (Fig. 2A and Fig. 2C). These results are compatible with the experimental observation. Given that the MCU has a low Ca^2+^ affinity, i.e., it requires high concentrations of Ca^2+^ for channel activation, increasing the mitochondrial surface portion that faces MAMs, and thus exposing more MCUs to higher concentrations of Ca^2+^, would lead to augmented MCU Ca^2+^ flux, and consequently, induce larger amplitudes of mitochondrial Ca^2+^ activities.

**Fig 2.**
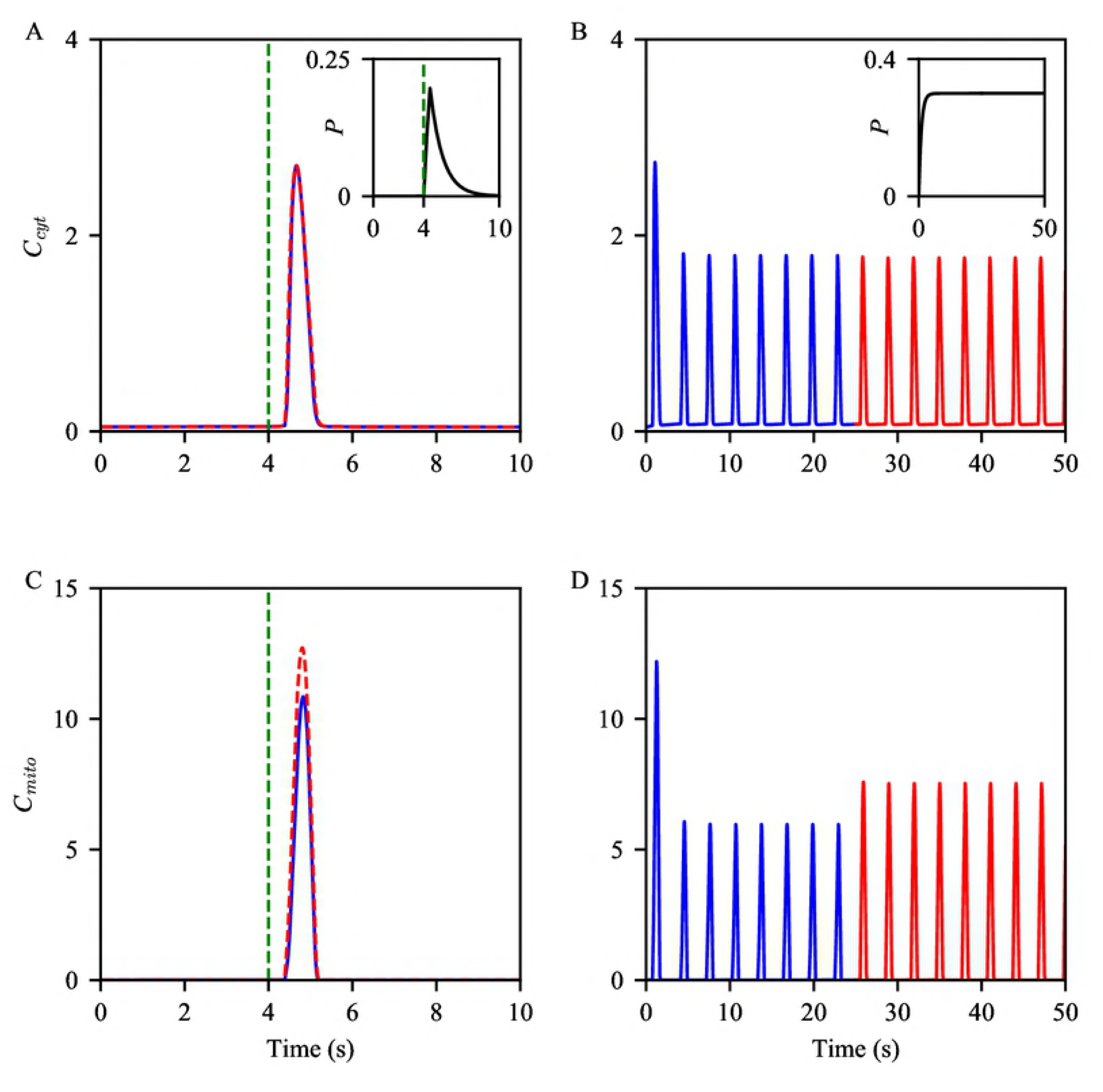
Effects of increased MAMs on amplitudes of cytosolic (top) and mitochondrial (bottom) Ca^2+^ activities. (A and C) We stimulated the model with a pulse of IP_3_, Eq. 2 with *M* = 0.5, *t*_0_ = 4, and ∆ = 0.5, shown by the inset graph in (A). The blue solid spikes were simulated with (*R*_S1_, *R*_S2_) = (0.2, 0.1), while the red dashed spikes were generated with the increased MAM surface ratios, (*R*_S1_, *R*_S2_) = (0.3, 0.15). The green dashed lines indicate the onset of the pulse. (B and D) The model was given continuous stimulation from IP_3_, which had a saturating concentration at *P*_s_ = 0.3, shown by the inset graph in (B). The blue oscillations were generated with the control parameters. Again, when we generated the red oscillations, *R*_S1_ and *R*_S2_ were increased by 50%.

We also studied model behavior under continuous stimulation. Instead of producing IP_3_ in a pulsatile manner, we modeled the concentration of IP_3_ to gradually increase until it reaches a steady state, *P*_s_. The model exhibited stable oscillations in cytosolic and mitochondrial Ca^2+^ concentrations (Fig. 2B and Fig. 2D). The increases of the *R*_S_ ratios were followed by an increased amplitude in mitochondrial oscillations, while the amplitude change in the cytosolic Ca^2+^ oscillations was negligible.

### Ca^2+^ activities in MAMs

Experimental observations suggest that Ca^2+^ concentration in MAMs is about 10-fold higher than that of the cytosol, and about 10-fold lower than that in the ER lumen [28, 29]. We examined whether the model reproduces these phenomena. The model was continuously stimulated with a constant IP_3_ concentration at its steady state *P*_s_ = 0.3 *µ*M to generate stable oscillations in all three compartments (Fig. 3A). The model simulated the expected order differences in the Ca^2+^ concentrations between the domains. We note that this model behavior is solely induced from the model assumption of a significantly large volume difference between the cytosol and MAMs. Since the portion of the ER membrane that faces the cytosol is larger than the other section juxtaposing MAMs, it is not surprising to see the larger cytosolic IPR Ca^2+^ flux (Fig. 3B and Fig. 3C). Nonetheless, the model simulated MAM Ca^2+^ oscillations with higher peaks, despite the relatively smaller MAM IPR Ca^2+^ fluxes.

**Fig 3.**
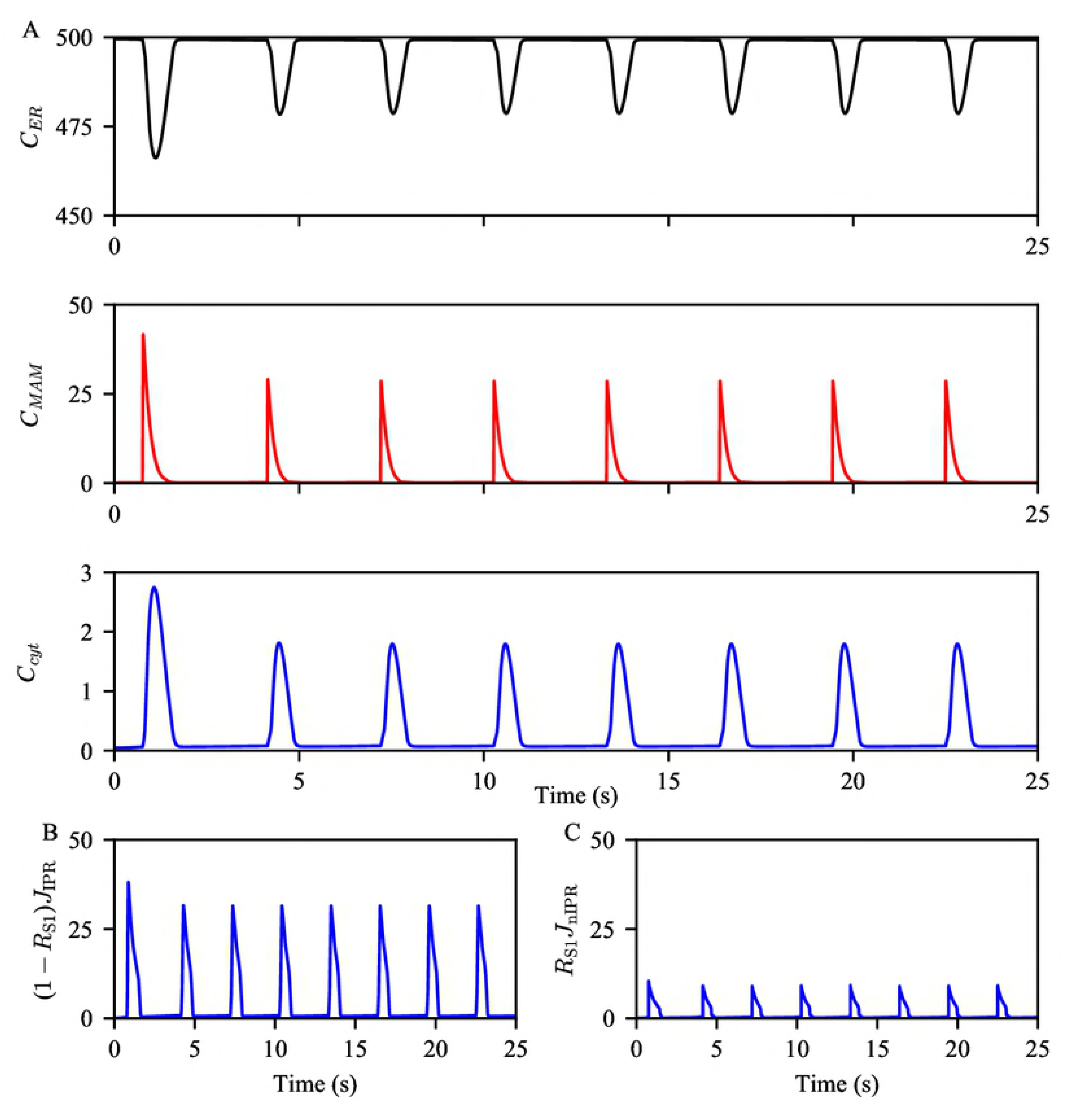
Ca^2+^ oscillations generated from the model exhibit varying orders of magnitude in different compartments. (A) The model was given continuous stimulation of IP_3_ with *P*_s_ = 0.3 *µ*M. From the top, the panels show Ca^2+^ oscillations in the ER, the MAM, and the bulk cytosol. (B and C) The magnitudes of IPR Ca^2+^ fluxes from the ER to the bulk cytosol and the MAM, respectively, during the oscillations shown in (A).

### Comparing Ca^2+^ oscillations: Control vs. obesity

Arruda et al. [9] reported the effects of obesity on the morphology of hepatic ER and mitochondria. Their study involved lean mice and two different groups of obese mice, one that had been on high-fat diet (HFD) for 16 weeks, and the other, genetically obese (*ob/ob*) mice. One of their main findings was that both groups of obese mice had a greater proportion of MAMs in liver cells. The authors also examined the expression levels of ER and mitochondrial proteins in liver lysates from the mice. Their western blot analysis showed that the obese mice had a higher expression of the IPR and PACS-2, an ER-mitochondrial tether protein. Interestingly, the expression level of MCU was higher in the *ob/ob* mice, while the difference between the lean and the HFD mice was negligible. This suggests that the change in the expression of MCU may not occur at the early stages of obesity, although it is likely to be associated with obesity in the long-term.

Another part of their study traced intracellular Ca^2+^ activities, both in the cytosol and mitochondrial lumen. Liver cells from lean and obese mice were stimulated with ATP to induce subsequent Ca^2+^ releases from the ER. The results showed higher peaks of mitochondrial Ca^2+^ concentration in cells from obese mice, and similar increases in cytosolic Ca^2+^ concentrations. From this, the authors speculated that the higher mitochondrial Ca^2+^ peaks observed under obesity are related to having more ER-mitochondrial interactions in the cells, and consequently, increased direct Ca^2+^ transport through MAMs.

We do not know the exact population of Ca^2+^ channels in mouse hepatocytes. However, the experimental data provide some guidelines for the relative expression levels between the control (lean) group and the obese group. The parameters shown in Table are control values, and when we simulated the model with this parameter set, which will be referred to as the *control model* from here on, model outcomes were regarded as the baseline behaviors for later comparison. Based on the experimental results discussed above, we modified some of the parameters to reflect cellular changes that are associated with obesity. Table 2 shows the parameter modifications. The model with the modified parameters will be referred to as the *obesity model*. We compared behaviors of the obesity model to those of the control model to confirm that the models correctly capture the gap between Ca^2+^ activities observed in hepatocytes from lean and obese mice.

**Table 1.** Parameter values of the model. The parameters with ^*^ are modified from their suggested values as in Wacquier et al. [22].

| Parameter | Value (Unit) | Description |
| --- | --- | --- |
| Parameters for the $\text{Ca}^{2+}$ and $\text{IP}_3$ concentration equations | | |
| $R_{V1}$ | 2000 | the volume ratio between the cytosol and MAM |
| $R_{V2}$ | 10 | the volume ratio between the cytosol and the ER |
| $R_{V3}$ | 15 | the volume ratio between the cytosol and mitochondria |
| $R_{S1}$ | 0.2 | the proportion of the ER membrane surface that adjoins the MAM |
| $R_{S2}$ | 0.1 | the proportion of mitochondrial membrane surface that adjoins the MAM |
| $C_t$ | 50 ($\mu\text{M}$) | the total intracellular $\text{Ca}^{2+}$ concentration |
| $D_c$ | 0.1 | intracellular $\text{Ca}^{2+}$ diffusion rate |
| $D_p$ | 1 | intracellular $\text{IP}_3$ diffusion rate |
| $k_{\text{leak}}$ | 0.001 ($\text{s}^{-1}$) | leak $\text{Ca}^{2+}$ flux coefficient |
| $k_{\text{IPR}}$ | 0.3 ($\text{s}^{-1}$) | coefficient of the IPR flux entering the bulk cytosol |
| $k_{\text{nIPR}}$ | 0.3 ($\text{s}^{-1}$) | coefficient of the IPR flux entering the MAM |
| $V_{\text{SERCA}}$ | $30 \text{ } (\mu\text{M s}^{-1})$ | coefficient of the SERCA flux from the bulk cytosol |
| $V_{\text{nSERCA}}$ | $20 \text{ } (\mu\text{M s}^{-1})$ | coefficient of the SERCA flux from the MAM |
| $K_{\text{SERCA}}$ | $0.35 \text{ } (\mu\text{M})$ | half-maximal activating cytosolic $\text{Ca}^{2+}$ concentration of SERCA |
| $K_{\text{e}}$ | $0.05 \text{ } (\mu\text{M})$ | dissociation constant of ATP from SERCA |
| $V_{\text{MCU}}$ | $0.00005 \text{ } (\mu\text{M s}^{-1})$ | coefficient of the MCU flux from the bulk cytosol |
| $V_{\text{nMCU}}$ | $0.00005 \text{ } (\mu\text{M s}^{-1})$ | coefficient of the MCU flux from the MAM |
| $V_{\text{NCX}}$ | $0.5 \text{ } (\mu\text{M s}^{-1})$ | coefficient of the NCX flux entering the bulk cytosol |
| $V_{\text{nNCX}}$ | $0.5 \text{ } (\mu\text{M s}^{-1})$ | coefficient of the NCX flux entering the MAM |
| $K_1$ | $19 \text{ } (\mu\text{M})$ | dissociation constant for $\text{Ca}^{2+}$ translocation by the MCU |
| $K_2$ | $0.38 \text{ } (\mu\text{M})$ | dissociation constant for MCU activation by $\text{Ca}^{2+}$ |
| $p_1$ | $0.1 \text{ } (\text{mV}^{-1})$ | coefficient of MCU activity dependence on voltage |
| $p_2$ | $0.016 \text{ } (\text{mV}^{-1})$ | coefficient of NCX activity dependence on voltage |
| Parameters for the IPR model equations |  |  |
| $q_{26}$ | $10500 \text{ } (\text{s}^{-1})$ | transition rate from $C_2$ to $O_6$ |
| $q_{62}$ | $4010 \text{ } (\text{s}^{-1})$ | transition rate from $O_6$ to $C_2$ |
| $L_{\text{IPR}}$ | $0.2 \text{ } (\text{s}^{-1})$ | the rate at which $h_{42}$ and $h_{\text{n}42}$ reach their equilibria, if the channel is in $C_4$ |
| $H_{\text{IPR}}$ | $10 \text{ } (\text{s}^{-1})$ | the rate at which $h_{42}$ and $h_{\text{n}42}$ reach their equilibria, if the channel is in the drive mode |
| $C_{p0}$ | $700 \text{ } (\mu\text{M})$ | $\text{Ca}^{2+}$ concentration at the pore of an open IPR |
| Parameters for the mitochondrial metabolic pathway equations |  |  |
| $N_{\text{mito}}^{\text{tot}}$ | $250 \text{ } (\mu\text{M})$ | the total mitochondrial pyridine nucleotide concentration |
| $A_{\text{mito}}^{\text{tot}}$ | $15000 \text{ } (\mu\text{M})$ | the total mitochondrial adenine nucleotide concentration |
| $A_{\text{cyt}}^{\text{tot}}$ | $2500 \text{ } (\mu\text{M})$ | the total cytosolic adenine nucleotide concentration |
| $L_{\text{MCU}}$ | 50 | allosteric equilibrium constant for uniporter conformations |
| $a_1^*$ | 10 | scaling factor between NADH oxidation and change in mitochondrial membrane voltage |
| $a_2$ | 3.43 | scaling factor between ADP phosphorylation and change in mitochondrial membrane voltage |
| $V_{\text{hyd}}^*$ | 150 ($\mu\text{M s}^{-1}$) | maximum rate of ATP hydrolysis |
| $K_{\text{hyd}}$ | 1000 ($\mu\text{M}$) | half-maximal activating cytosolic ATP concentration of ATP hydrolysis |
| $V_{\text{ant}}$ | 5000 ($\mu\text{M s}^{-1}$) | rate coefficient of the adenine nucleotide translocator |
| $\alpha_c$ | 0.111 | cytosolic ADP and ATP buffering coefficient |
| $\alpha_m$ | 0.139 | mitochondrial ADP and ATP buffering coefficient |
| $F$ | 96480 ($\text{C mol}^{-1}$) | Faraday constant |
| $R$ | 8315 ($\text{mJ mol}^{-1} \text{K}^{-1}$) | perfect gas constant |
| $T$ | 310.16 (K) | temperature |
| $V_{\text{F}_1\text{F}_\text{O}}$ | 35000 ($\mu\text{M s}^{-1}$) | rate coefficient of the $\text{F}_1\text{F}_\text{O}$-ATPase |
| $k_{\text{gly}}$ | 450 ($\mu\text{M s}^{-1}$) | rate coefficient of glycolysis |
| $k_{\text{o}}$ | 600 ($\mu\text{M s}^{-1}$) | rate coefficient of NADH oxidation by ETC |
| $V_{\text{agc}}^*$ | 180 ($\mu\text{M s}^{-1}$) | rate coefficient of NADH production |
| $K_{\text{agc}}$ | 0.14 ($\mu\text{M}$) | dissociation constant of cytosolic $\text{Ca}^{2+}$ from AGC |
| $p_4$ | 0.01 ($\text{mV}^{-1}$) | coefficient of AGC activity dependence on voltage |
| $q_1$ | 1 | Michaelis-Menten-like constant for $\text{NAD}^+$ consumption by the TCA |
| $q_2$ | 0.1 ($\mu\text{M}$) | half-maximal activating mitochondrial $\text{Ca}^{2+}$ concentration of the TCA |
| $q_3$ | 100 (mV) | Michaelis-Menten constant for NADH consumption by the ETC |
| $q_4$ | 177 (mV) | coefficient of ETC activity dependence on voltage |
| $q_5$ | 5 (mV) | coefficient of ETC activity dependence on voltage |
| $q_6$ | 10000 ($\mu\text{M}$) | inhibition constant of ATPase activity by ATP |
| $q_7$ | 190 (mV) | coefficient of ATPase activity dependence on voltage |
| $q_8$ | 8.5 (mV) | coefficient of ATPase activity dependence on voltage |
| $q_9$ | $2 \text{ } (\mu\text{M s}^{-1} \text{ mV}^{-1})$ | proton leak dependence on voltage |
| $q_{10}$ | $-30 \text{ } (\mu\text{M s}^{-1})$ | rate coefficient of voltage-independent proton leak |

**Table 2.** Modified parameters for the obesity model simulations.

| Parameter | control model | obesity model |
| --- | --- | --- |
| $R_{S1}$ | 0.2 | 0.3 |
| $R_{S2}$ | 0.1 | 0.15 |
| $k_{\text{IPR}}$ | $0.3 \text{ s}^{-1}$ | $0.36 \text{ s}^{-1}$ |
| $k_{\text{nIPR}}$ | $0.3 \text{ s}^{-1}$ | $0.36 \text{ s}^{-1}$ |
| $V_{\text{MCU}}$ | $0.00005 \text{ } \mu\text{M s}^{-1}$ | $0.0001 \text{ } \mu\text{M s}^{-1}$ |
| $V_{\text{nMCU}}$ | $0.00005 \text{ } \mu\text{M s}^{-1}$ | $0.0001 \text{ } \mu\text{M s}^{-1}$ |

The panels in Fig. 4 show Ca^2+^ oscillations generated from the control and obesity models with the same magnitude of stimulation. The simulations suggest that hepatocytes from obese mice may exhibit faster Ca^2+^ oscillations with a higher amplitude of mitochondrial Ca^2+^, compared to cells from lean mice. In fact, the amplitude in the cytosol showed a relatively small change (*≈* 5%), compared to the mitochondrial Ca^2+^ amplitude, which was increased by 126%. From the experimental data, we cannot distinguish which cellular changes are responsible for which effects, and to what degree. Using our Ca^2+^ model, we investigated how Ca^2+^ oscillations are modulated by each cellular change associated with obesity. Model results are discussed in the following subsections.

**Fig 4.**
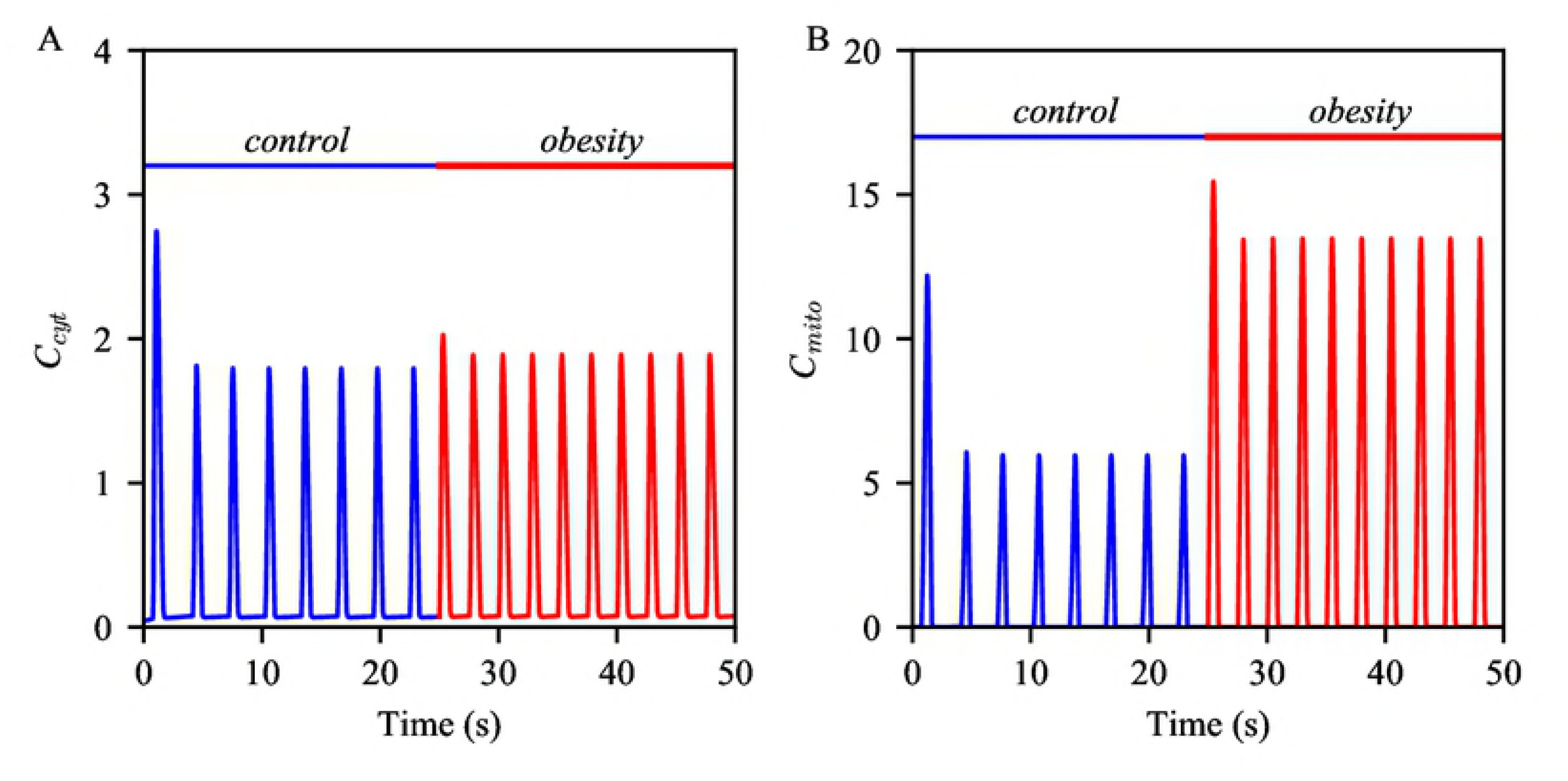
Effects of the cellular changes associated with obesity on Ca^2+^ oscillations. Oscillations of (A) cytosolic and (B) mitochondrial Ca^2+^ concentrations. The blue oscillations were generated from the model with the control parameters. When simulating the red oscillations, some of the parameters were modified as in Table 2. Throughout the simulation, IP_3_ concentration was held constant at 0.3 *µ*M.

### Increased MAM formation

We have previously discussed that a possible mechanism underlying the amplitude change in mitochondrial Ca^2+^ activities associated with the enhancement of MAM formation is a synergistic process that comes from the MCU’s low Ca^2+^ affinity and the small volume of MAMs. The increase in the amplitude of mitochondrial Ca^2+^ oscillations induced from the increased degree of MAM formation is already evident in Fig. 2. Interestingly, expressing more MAMs in the model slightly increased the oscillation frequency. Table 3 shows the quantified effects of 20%, 50% and 100% increase in MAM formation on the amplitudes and frequency of cytosolic and mitochondrial Ca^2+^ oscillations. The parameter adjustments induced a trend of negative effects on the amplitude of cytosolic Ca^2+^ oscillations, while having positive effects on that of mitochondrial oscillations. As stated in the introduction, cells tightly regulate frequencies and amplitudes of Ca^2+^ oscillations to orchestrate many cellular activities. Thus, although the magnitude of the frequency change is small, its effect on cellular homeostasis may be significant.

**Table 3.** Effects of increasing the degree of MAM formation, IPR activity, and MCU activity on the amplitudes of cytosolic and mitochondrial Ca^2+^ oscillations and the oscillation frequency.

| $\Delta$ in MAMs | $\Delta$ in $C_{cyt}$ amplitude | $\Delta$ in $C_{mito}$ amplitude | $\Delta$ in oscillation freq. |
| --- | --- | --- | --- |
| 20% | -0.56% | 10% | 0.5% |
| 50% | -1.3% | 26% | 0.8% |
| 100% | -2.4% | 55% | 0.8% |
| $\Delta$ in $k_{IPR}$, $k_{nIPR}$ | | | |
| 20% | 17% | 42% | 30% |
| 50% | 39% | 99% | 61% |
| 100% | 68% | 181% | 84% |
| $\Delta$ in $V_{MCU}$, $V_{nMCU}$ | | | |
| 20% | -1.4% | 14% | -0.7% |
| 50% | -3.2% | 33% | -1.6% |
| 100% | -6% | 61% | -2.9% |
| 200% | -11% | 104% | -4.4% |

### Increased IPR activity

Secondly, we looked at the effects of increased IPR activity on Ca^2+^ oscillations, by increasing *k*_IPR_ and *k*_nIPR_, the rate constants for the IPR Ca^2+^ fluxes. These adjustments lead to an increase in the magnitude of total Ca^2+^ flux through the IPR, either from an increased number of channels or an increase in the maximum strength of each channel. Since the model is a deterministic model that incorporates all-or-none activation of IPRs, we intuitively expected the increased IPR parameters to generate Ca^2+^ oscillations with larger amplitudes in both the cytosol and mitochondria. The model was stimulated with *P*_s_ = 0.3 *µ*M to produce the stable oscillations in Fig. 5. At *t* = 25 s, only *k*_IPR_ and *k*_nIPR_ were increased by 20%, while other parameters were kept the same. The parameter adjustments induced increases in the amplitudes of cytosolic and mitochondrial Ca^2+^ oscillations by 17% and 42%, respectively, as expected. The model simulations also showed a 30% increase in the oscillation frequency. With the increased IPR channel activity, it takes less time for cytosolic Ca^2+^ concentration to reach the threshold concentration for spiking. However, this also means that the channels have a shorter refractory time, the period of time that it takes for the IPR to recover its state between spikes, and hence, at the time of spiking, the receptors would have a lower open probability. Fig. 5C and Fig. 5D confirm this conjecture.

**Fig 5.**
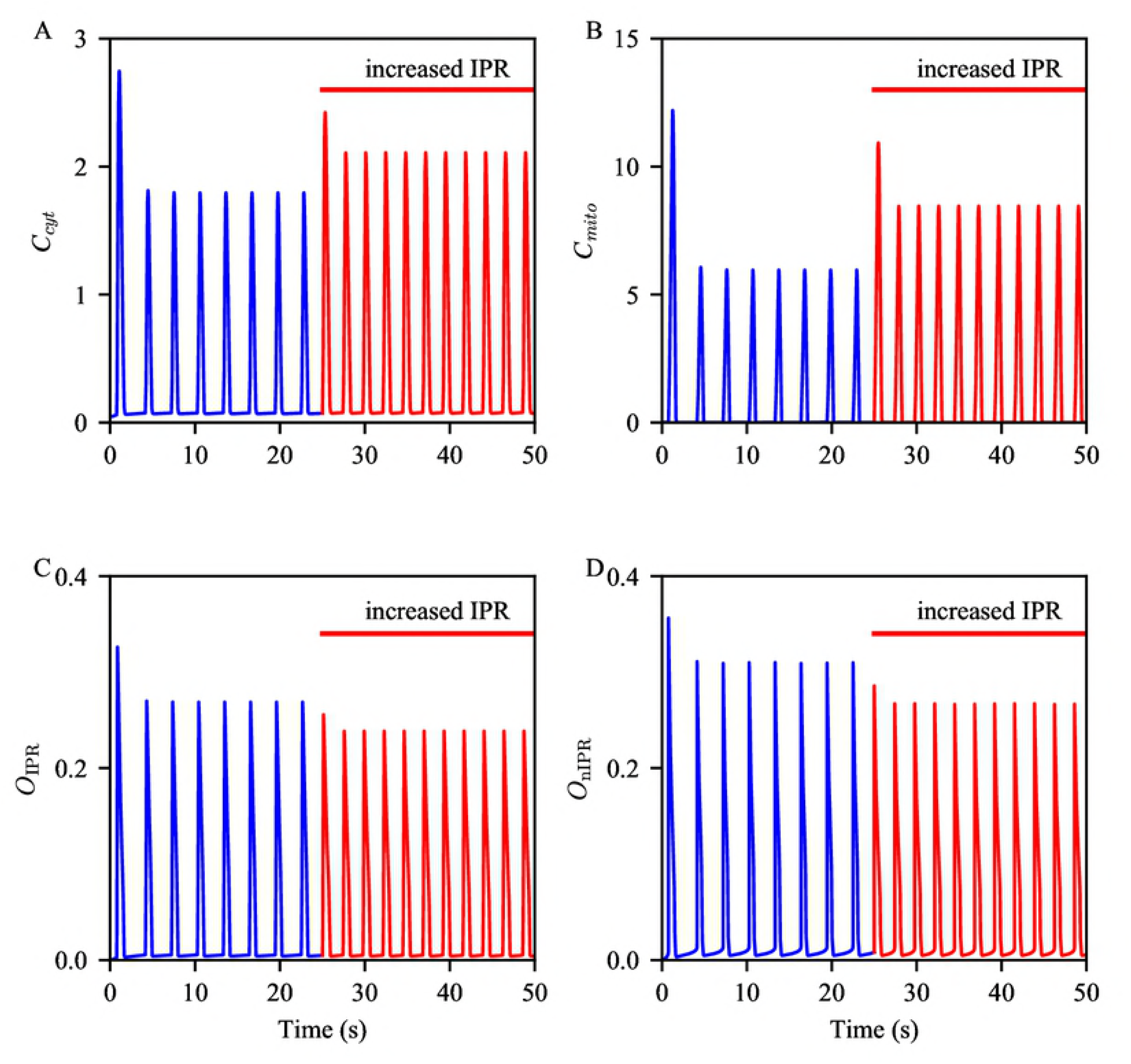
Effects of increased IPR activity on Ca^2+^ oscillations and IPR open probabilities. (A and B) Oscillations in cytosolic and mitochondrial Ca^2+^ concentrations, respectively. The blue oscillations are generated from the control model, while the red oscillations are generated with increased *k*_IPR_ and *k*_nIPR_. The model was given continuous stimulation of IP_3_ with *P*_s_ = 0.3 *µ*M. (C and D) Corresponding open probabilities of the bulk cytosol IPR and the MAM IPR, respectively.

We further increased *k*_IPR_ and *k*_nIPR_ by 50%, and then by 100%, to affirm the positive correlation between the parameters and the oscillation amplitudes and frequency (Table 3). It is evident that the corresponding parameter modifications had strong positive effects on all three properties of oscillations that were of interest.

### Increased MCU activity

Lastly, we simulated the effects of an increase in MCU activity level on cytosolic and mitochondrial Ca^2+^ oscillations. Following the same steps as the previous simulations, the model was given continuous stimulation with *P*_s_ = 0.3 *µ*M, which gave rise to the stable oscillations shown in the panels of Fig. 6. At *t* = 25 s, *V*_MCU_ and *V*_nMCU_ were increased by 100%, and the model continued to exhibit oscillations shown in the figures. The increased parameters had a negative effect on the amplitude of cytosolic Ca^2+^ oscillations and the oscillation frequency, whereas the amplitude of mitochondrial Ca^2+^ oscillations was increased. The relative magnitudes of these changes are quantified in Table 3. The most obvious explanation for the amplitude changes is that the enhanced activity of MCUs transports a larger amount of Ca^2+^ from the cytosol to mitochondria, and the magnitude of change is more substantial in mitochondria than in the cytosol due to the compartments’ volume difference. We also examined the changes under 20%, 50%, and 200% increase of the parameters, to verify the overall trend of the effects.

**Fig 6.**
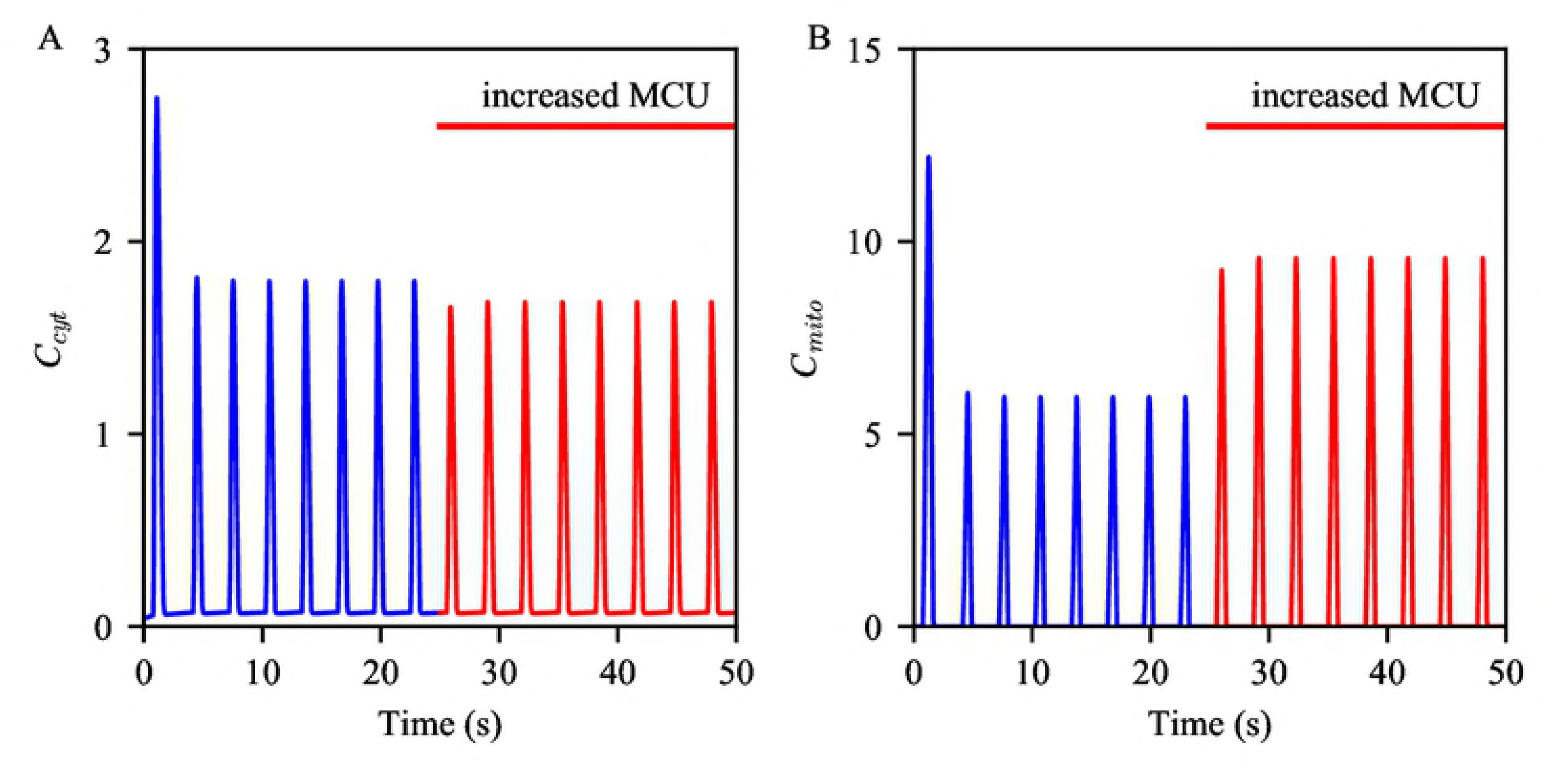
Effects of increased MCU activity on Ca^2+^ oscillations. Oscillations of (A) cytosolic and (B) mitochondrial Ca^2+^ concentrations generated from the control model, in blue, and with increased *V*_MCU_ and *V*_nMCU_, in red. The model was given continuous stimulation of IP_3_ with *P*_s_ = 0.3 *µ*M.

As mentioned above, no significant difference was reported between the expression levels of MCU in liver cells from the lean and the HFD mice. However, cells from *ob/ob* mice had a marked increase in MCU expression level. These experimental data suggest that the increase in the MCU expression level may not be a direct response to the onset of obesity. We have shown that both increases of MAM formation and IPR channel density, the cellular changes that were observed in the HFD mice, and thus are early events associated with obesity, induce higher peaks in mitochondrial Ca^2+^.

### Model prediction

We postulate that liver cells from different health conditions have varying thresholds for the breaking point of their Ca^2+^ oscillations. In order to test this, we studied model behaviors under three different sets of parameters: one is without any modification, which represents the healthy case, another one is with increased *R*_S_’s, *k*_IPR_ and *k*_nIPR_, which corresponds to the cell condition associated with HFD, and the last set is with additional parameter increases of *V*_MCU_ and *V*_nMCU_, which refers to cells from *ob/ob* animals. For each parameter set, the model was given continuous stimulation with five distinct regimes where *P*_s_ is incrementally adjusted, as shown in the panels of Fig. 7. It is clearly shown that the model representing the *ob/ob* condition reached the breaking point for Ca^2+^ oscillations at a lower concentration of IP_3_, compared to the HFD and healthy conditions. The control model exhibited Ca^2+^ oscillations that are robust to higher concentrations of IP_3_.

**Fig 7.**
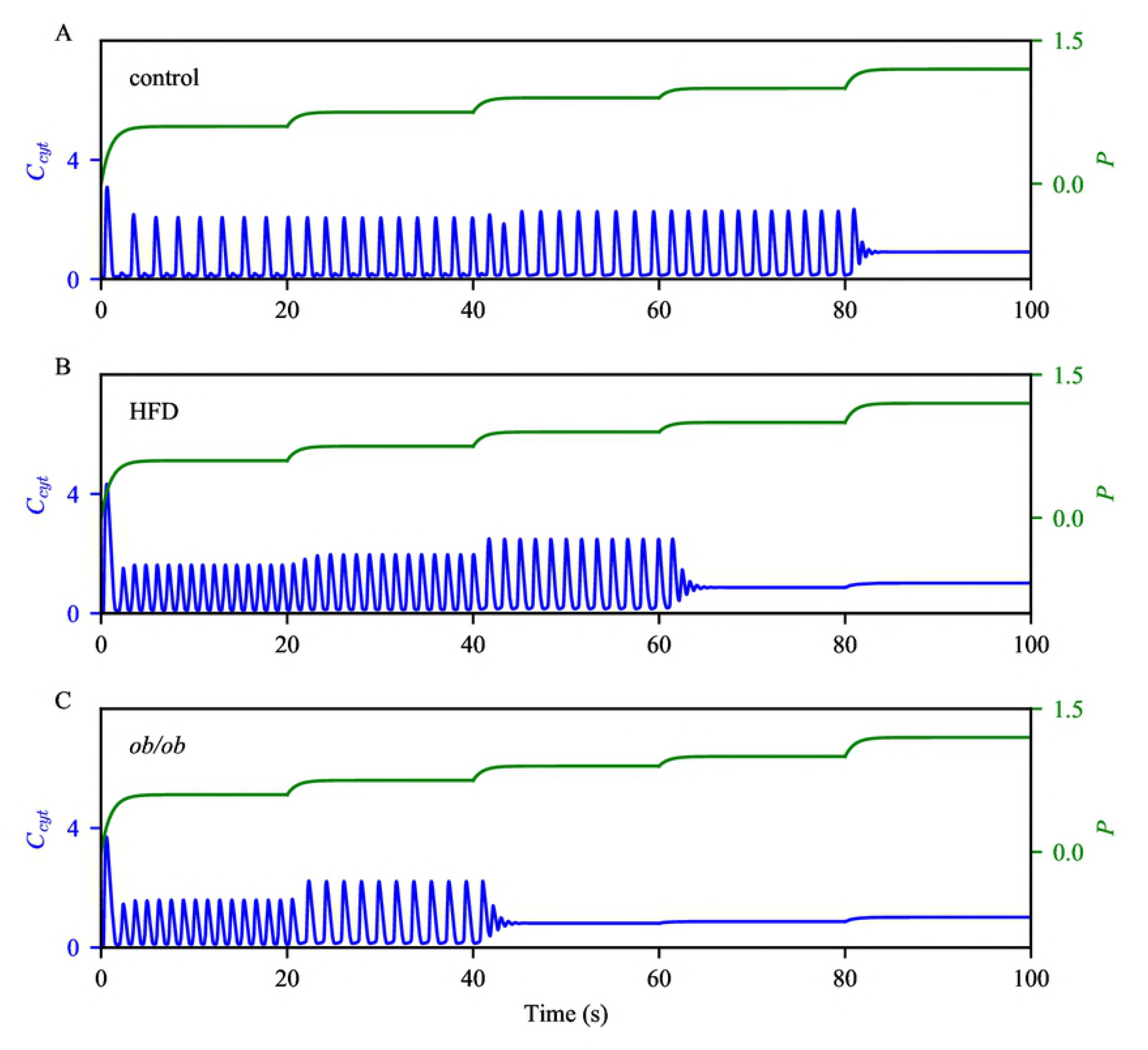
Robustness of Ca^2+^ oscillations under different model conditions. We perturbed the model with gradually increasing stimulation, under three model conditions: (A) control, (B) HFD, and (C) *ob/ob*. Initially, *P*_s_ was at 0.6 *µ*M, then was increased to 0.75 *µ*M, 0.9 *µ*M, 1 *µ*M, and then to 1.2 *µ*M at *t* = 20 s, 40 s, 60 s, and 80 s, respectively. The cytosolic Ca^2+^ concentrations are shown in blue, with the scale on the left y-axis. The green timeseries represent the cytosolic IP_3_ concentration, with the scale on the right y-axis.

To verify these predictions, we suggest measuring Ca^2+^ responses in liver cells with each of the three conditions to a wide range of IP_3_ concentration, and see if the average IP_3_ oscillatory range was decreased in the *ob/ob* condition. However, one of the challenges of this experiment is that, due to cell-to-cell variability, it is critical to measure responses from a myriad of cells before drawing a conclusion.

## Discussion

We have presented a mathematical model for Ca^2+^ dynamics in mouse hepatocytes. Based on experimental data that showed oscillatory Ca^2+^ signals in hepatocytes in Ca^2+^-free media, the model is the closed-cell type, wherein the total intracellular Ca^2+^ concentration is assumed to be conserved (i.e., there is no Ca^2+^ flux across the plasma membrane). To our knowledge, this model is the first mathematical model to explicitly express Ca^2+^ concentration in MAMs as a dynamical variable and also show MAM Ca^2+^ levels to be within reasonable proportions of the Ca^2+^ levels in the other domains. The first aim of the model was to reproduced the data reported by Arruda et al. where hepatocytes with more MAMs exhibited ATP-induced Ca^2+^ transient with higher peaks in mitochondrial Ca^2+^ concentration [9]. We assumed that the IPRs, MCUs and NCXs are uniformly expressed on the residing membranes, while the SERCA pumps are predominant on the cytosolic side of the ER membrane.

Arruda et al. [9] also compared hepatic cellular characteristics between different groups of mice. They had a group of lean mice as their control, and two groups of mice, one that had been under high fat diet (HFD) and the other genetically obese (*ob/ob*) mice, for mouse models of obesity. The *ob/ob* mouse cells showed higher expression levels of IPR and MCU, as well as a higher degree of MAM formation. The HFD mouse cells also had more MAMs and a higher IPR channel density than the lean group, but the groups’ MCU expression levels did not differ. We interpreted the morphological change of MAMs and and the increase in IPR channel density as the traits of obesity in early stages, while considering the increase in MCU channel density as an effect that appears in later stages.

We used the model to study how Ca^2+^ signals are altered by each cellular change associated with obesity. According to model simulations, an increase of MAM proportion speeds up Ca^2+^ oscillations and increases mitochondrial Ca^2+^ amplitudes, while decreasing the amplitude of cytosolic Ca^2+^. An increase in IPR channel activity level induces faster Ca^2+^ oscillations with higher amplitudes in both cytosol and mitochondria. Furthermore, an increase in the level of MCU activity increases mitochondrial Ca^2+^ amplitude and oscillation period. We found that the cellular changes observed in the HFD mouse cells have a positive effect on oscillation frequency, while the additional change that only appeared in the *ob/ob* has the opposite effect. Thus, we postulate that the increase of MCU channel density observed in the *ob/ob* mouse cells is a secondary effect of obesity that counteracts other cellular changes that occur in the HFD mouse cells.

Metabolic flexibility refers to the ability of the organism to adapt its fuel source, depending on availability and need [30], and emerging evidence suggests the involvement of MAMs in metabolic flexibility [31]. Interestingly, Rieusset et al. [32] reported a link between the disruption of ER-mitochondrial Ca^2+^ exchange and hepatic insulin resistance in their mouse model. As we have shown in this paper, hepatic cellular changes associated with HFD and obesity affect Ca^2+^ oscillation frequencies and amplitudes. However, the question of whether the altered Ca^2+^ dynamics plays a causal role in the development of hepatic insulin resistance and metabolic diseases remains to be explored. We eagerly anticipate that our model can be one of stepping stones addressing such conundrum.

### Mitochondrial Ca^2+^ oscillations and ROS

There are many studies that associate mitochondrial Ca^2+^ dynamics with the generation of ROS [8, 33, 34]. It has been suggested that mitochondrial Ca^2+^ up-regulates ETC, which in turn increases the generation of mitochondrial superoxide, one of the main forms of ROS. As a counter mechanism, Ca^2+^ also activates enzymes that scavenge superoxide. The optimal feedback mechanisms between mitochondrial Ca^2+^ and ROS would keep ROS at a tolerable level to avoid ROS-induced apoptosis. Achieving this state requires a balance between the production and degradation of ROS, and most likely in a time-delay manner. We have shown that the cellular changes associated with HFD induce faster Ca^2+^ oscillations in mitochondria, which potentially leads to a faster rate of ROS production. If the process of ROS degradation cannot keep up with the increased level of ROS, it would cause the accumulation of ROS in mitochondria and consequently, apoptosis. However, according to the model simulations, oscillation frequencies can be decreased by an increase of MCU channel density, another cellular change associated with obesity. Unfortunately, this reaction potentially leads to mitochondrial Ca^2+^ overload, which is a deleterious phenomenon for cells. There are many studies that discuss links between mitochondrial Ca^2+^ overload and oxidative stress and ER stress, which are the hallmark of apoptosis [3, 35–38]. Nonetheless, liver cells from the *ob/ob* mice have an increased expression level of MCU, which aggravates mitochondrial Ca^2+^ overload. The fact that the cells increase their MCU channels on top of the other cellular changes is an interesting and physiologically counterintuitive behavior. Why would they push themselves in an adverse direction and worsen their condition? Based on our model simulations, we conjecture that the MCU expression level change is a counter response to the other cellular changes. The increases in MAM formation and IPR channel density both accelerated the oscillations, while the increase in MCU channel density had an opposite effect on the oscillations. Thus, we suggest that cells increase their MCU expression level in an attempt to adjust their oscillation frequency to a sustainable level. Of course, this mechanism has to be finely controlled, as there are other problems, such as mitochondrial Ca^2+^ overload, that come with it.

### Other Ca^2+^ fluxes

As discussed before, we presented a closed-cell model, and did not include Ca^2+^ fluxes across the plasma membrane, such as the ones via store-operated Ca^2+^ channels (SOCCs), receptor-operated Ca^2+^ channels (ROCCs), and plasma membrane Ca^2+^-ATPase (PMCA) pumps. SOCCs draw Ca^2+^ from the extracellular space when Ca^2+^ stores (ER/SR) get depleted. ROCCs, which also carry Ca^2+^ into the cytosol, are activated by agonists binding on G-protein-coupled receptors. PMCA pumps remove Ca^2+^ from the cytosol. The fluxes through these channels also modulate intracellular Ca^2+^ dynamic, and thus could be added in the model. In particular, experimental evidence suggests that mitochondrial Ca^2+^ regulate SOCCs [39–41]. When mitochondria sequester a substantial amount of Ca^2+^ from the cytosol and the ER, the level of Ca^2+^ concentration in the ER could get sufficiently low and activate SOCCs.

The model proposed by Wacquier et al. [22] includes a bidirectional Ca^2+^ flux between the cytosol and mitochondria. They suggested that this flux could represent the mitochondrial permeability transition pore (PTP) in the low conductance mode. It would be interesting to see how our model behaves in the presence of the bidirectional flux, and compare the models.

### Model reduction

Due to the high dimensionality of the model, which is a system of 11 ODEs, numerical integration can be computationally expensive. In an attempt to simplify the model, we applied quasi-steady state reduction and assumed that the activation variables of the IPRs, *h*_42_ and *h_n_*_42_, instantaneously reached their quasi-equilibria. However, this reduction destroyed the desired order differences between the magnitudes of Ca^2+^ transients in the cytosol, the ER, and MAMs. This implies that the kinetics of IPR activation is an important factor for modeling the 10-fold difference in the amplitudes of cytosolic and MAMs Ca^2+^ activities, shown in Fig. 3.

## Supporting information

**S1 .ode file.** The model’s .ode file that runs on XPPAUT [27].

## Acknowledgments

We thank James Sneyd at University of Auckland, New Zealand, and Arthur Sherman, our lab chief, for their helpful discussions during the course of this work.

## References

1. Berridge MJ, Lipp P, Bootman MD. The versatility and universality of calcium signalling. Nat Rev Mol Cell Biol. 2000;1:11–21.

2. Arruda AP, Hotamisligil GS. Calcium homeostasis and organelle function in the pathogenesis of obesity and diabetes. Cell Metab. 2015;22(3):381–397.

3. Gӧrlach A, Bertram K, Hudecova S, Krizanova O. Calcium and ROS: A mutual interplay. Redox Biol. 2015;6:260–271.

4. Patergnani S, et al. Calcium signaling around Mitochondria associated membranes (MAMs). Cell Commun Signal. 2011;9:19.

5. Cárdenas-Pérez RE, Camancho A. Roles of calcium and mitochondria-associated membranes in the development of obesity and diabetes. Medicina Universitaria. 2016;18(70):23–33.

6. Mekahli D, Bultynck G, Parys JB, De Smedt H, Missiaen L. Endoplasmic-reticulum calcium depletion and disease. Cold Spring Harb Perspect Biol. 2011;3(6):a004317. doi:10.1101/cshperspect.a004317.

7. Krols M, van Isterdael G, Asselbergh B, Kremer A, Lippens S, Timmerman V, et al. Mitochondria-associated membranes as hubs for neurodegeneration. Acta Neuropathol. 2016;131:505–523. doi:10.1007/s00401-015-1528-7.

8. Brookes PS, Yoon Y, Robotham JL, Anders MW, Sheu SS. Calcium, ATP, and ROS: a mitochondrial love-hate triangle. Am J Physiol, Cell Physiol. 2004;287:C817–C833. doi:10.1152/ajpcell.00139.2004.

9. Arruda AP, Pers BM, Parlakgül G, Güney E, Inouye K, Hotamisligil GS. Chronic enrichment of hepatic endoplasmic reticulum-mitochondria contact leads to mitochondrial dysfunction in obesity. Nat Med. 2014;20(12):1427–1435.

10. Li YX, Rinzel J. Equations for InsP_3_ receptor-mediated [Ca^2+^] oscillations derived from a detailed kinetic model: a Hodgkin-Huxley like formalism. J Theor Biol. 1994;166:461–473.

11. De Young GW, Keizer J. A single-pool inositol 1,4,5-trisphosphate-receptor-based model for agonist-stimulated oscillations in Ca^2+^ concentration. Proc Natl Acad Sci USA. 1992;89:9895–9899.

12. Shuai JW, Jung P. Stochastic properties of Ca^2+^ release of inositol 1,4,5-trisphosphate receptor clusters. Biophys J. 2002;87(1):87–97.

13. Dupont G, Goldbeter A. One-pool model for Ca^2+^ oscillations involving Ca^2+^ and inositol 1,4,5-trisphosphate as co-agonist for Ca^2+^ release. Cell Calcium. 1993;14:311–322.

14. Cortassa S, Aon MA, Marbán E, Winslow RL, O’Rouke B. An integrated model of cardiac mitochondrial energy metabolism and calcium dynamics. Biophys J. 2003;84(4):2734–2755.

15. Bertram R, Gram Pedersen M, Luciani DS, Sherman A. A simplified model for mitochondrial ATP production. J Theor Biol. 2006;243:575–586.

16. Magnus G, Keizer J. Minimal model of β-cell mitochondrial Ca^2+^ handling. Am J Physiol. 1997;273 (2 Pt 1):C717–C733.

17. Nguyen MHT, Jafri MS. Mitochondrial calcium signaling and energy metabolism. Ann N Y Acad Sci. 2005;1047:127–137. doi:10.1196/annals.1341.012.

18. Fall CP, Keizer JE. Mitochondrial modulation of intracellular Ca^2+^ signaling. J Theor Biol. 2001;210:151–165.

19. Patterson M, Sneyd J, Friel DD. Depolarization-induced calcium responses in Sympathetic Neurons: Relative Contributions from Ca^2+^ Entry, Extrusion, ER/Mitochondrial Ca^2+^ Uptake and Release, and Ca^2+^ Buffering. J Gen Physiol. 2007;129(1):29–56. doi:10.1085/jgp.200609660.

20. Szopa P, Dyzma M, Kázmierczak B. Membrane associated complexes in calcium dynamics modelling. Phys Biol. 2013;10(3):035004.

21. Qi H, Li L, Shuai J. Optimal microdomain crosstalk between endoplasmic reticulum and mitochondria for Ca^2+^ oscillations. Sci Rep. 2015;5:7984. doi:10.1038/srep07984.

22. Wacquier B, Combettes L, Tran Van Nhieu G, Dupont G. Interplay between intracellular Ca^2+^ oscillations and Ca^2+^-stimulated mitochondrial metabolism. Sci Rep. 2016;6:19316. doi:10.1038/srep19316.

23. Penny CJ, Kilpatrick BS, Han JM, Sneyd J, Patel S. A computational model of lysosome-ER Ca^2+^ microdomains. J Cell Sci. 2014;127:2934–2943.

24. Jones BF, Boyles RR, Hwang S, Bird GS, Putney JW. Calcium influx mechanisms underlying calcium oscillations in rat hepatocytes. Hepatology. 2008;48(4):1273–1281.

25. Cao P, Tan X, Donovan G, Sanderson MJ, Sneyd J. A deterministic model predicts the properties of stochastic calcium oscillations in airway smooth muscle cells. PLoS Comput Biol. 2014;10(8):e1003783.

26. Hajnóczky G, Thomas AP. Minimal requirements for calcium oscillations driven by the IP_3_ receptor. EMBO J. 1997;16(12):3533–3543.

27. Ermentrout B. Simulating, analyzing, and animating dynamical systems: A guide to XPPAUT for researchers and students. SIAM Reviews. 2003;45(1):150–152.

28. Giacomello M, Drago I, Bortolozzi M, Scorzeto M, Gianelle A, Pizzo P, et al. Ca^2+^ Hot Spots on the Mitochondrial Surface AreGenerated by Ca^2+^ Mobilization from Stores, but Not by Activation of Store-Operated Ca^2+^ Channels. Mol Cel. 2010;38(2):280–290. doi:10.1016/j.molcel.2010.04.003.

29. Tang S, Wong HC, Wang ZM, Huang Y, Zou J, Zhuo Y, et al. Design and application of a class of sensors to monitor Ca^2+^ dynamics in high Ca^2+^ concentration cellular compartments. Proc Natl Acad Sci USA. 2011;108(39):16265–16270. doi:10.1073/pnas.1103015108.

30. Galgani JE, Moro C, Ravussin E. Metabolic flexibility and insulin resistance. Am J Physiol Endocrinol Metab. 2008;295(5):E1009–E1017. doi:10.1152/ajpendo.90558.2008.

31. Theurey P, Rieusset J. Mitochondria-associated membranes response to nutrient availability and role in metabolic diseases. Trends in Endocrinology & Metabolism. 2017;28(1):32–45. doi:https://doi.org/10.1016/j.tem.2016.09.002.

32. Rieusset J, Fauconnier J, Paillard M, Belaidi E, Tubbs E, Chauvin M, et al. Disruption of calcium transfer from ER to mitochondria links alterations of mitochondria-associated ER membrane integrity to hepatic insulin resistance. Diabetologia. 2016;59(3):614–623. doi:https://doi.org/10.1007/s00125-015-3829-8.

33. Adam-Vizi V, Starkov AA. Calcium and Mitochondrial Reactive Oxygen Species Generation: How to Read the Facts. J Alzheimers Dis. 2010;20:S413–S426. doi:10.3233/JAD-2010-100465.

34. Csordás G, Hajnóczky G. SR/ER-mitochondrial local communication: Calcium and ROS. Biochim Biophys Acta. 2009;1787:1352–1362. doi:10.1016/j.bbabio.2009.06.004.

35. Hajnóczky G, Davies E, Madesh M. Calcium signaling and apoptosis. Biochem Biophys Res Commun. 2003;304(3):445–454. doi:10.1016/S0006-291X(03)00616-8.

36. Pinton P, Ferrari D, Rapizza E, Di Virgilio F, Pozzan T, Rizzuto R. A role for calcium in Bcl-2 action? Biochimie. 2002;84(2–3):195–201. doi:10.1016/S0300-9084(02)01373-1.

37. Pinton P, Giorgi C, Siviero R, Zecchini E, Rizzuto R. Calcium and apoptosis: ER-mitochondria Ca^2+^ transfer in the control of apoptosis. Oncogene. 2008;27(50):6407–6418. doi:10.1038/onc.2008.308.

38. Malhotra JD, Kaufman RJ. ER stress and its functional link to mitochondria: role in cell survival and death. Cold Spring Harb Perspect Biol. 2011;3:a004424. doi:10.1101/cshperspect.a004424.

39. Hoth M, Fanger CM, Lewis RS. Mitochondrial regulation of store-operated calcium signaling in T lymphocytes. J Cell Biol. 1997;137(3):633. doi:10.1083/jcb.137.3.633.

40. Glitsch MD, Bakowski D, Parekh AB. Store-operated Ca^2+^ entry depends on mitochondrial Ca^2+^ uptake. EMBO J. 2002;21(24):6744–6754.

41. Parekh AB, Putney JW. Store-operated calcium channels. Physiol Rev. 2005;85(2):757–810. doi:10.1152/physrev.00057.2003.

